# Identification of Blood Based Bio-Marker for Huntington’s disease using in-silico gene expression analysis

**DOI:** 10.1101/2021.03.26.437248

**Authors:** Souvik Chakraborty

## Abstract

Huntington’s disease (HD) is an autosomal dominant neurodegenerative disorder with profound phenotypic characters. HD is at present incurable and there are several trials going on to find a cure. HD is caused when there is a mutation in the Huntingtin gene which is found to be associated with axonal transport. Diagnosis is based on the signs and symptoms of the patients but by that time the psychomotor problems have already reached the level from where reversing the disease is impossible. Blood based biomarkers can be used for the diagnosis of the disease at an early stage. In this study several gene expression study data were analyzed and there were 329 Differentially Expressed Genes (DEGs) in all the three chosen datasets. Protein protein interaction network was created using STRING and CytoHubba plug-in was used to identify top ten hub genes which are CXCL8, PSMC6, UBE2D1, UBE2D1, CD27, UBE2D3, SF3B1, CASP3, EIF4E, BIRC2 and PTEN. Online software Enrichr was used for Gene Ontology and KEGG pathway enrichment analysis to find out the biological process, molecular function, cellular component and the pathways that were enriched in HD. This study finds out that those genes which were present in all the three datasets namely FNDC3A, BCLAF1 and ALCAM were not the hub genes. So further studies are required for identifying a potential biomarker of HD.

## Introduction

Huntington’s disease (HD) is a neurodegenerative disease which primarily affects motor system of our body and is associated with many phenotypic characteristics such as athetosis, chorea etc.^1^ HD affects approximately 1/20,000 people and the average age at the onset of the disease ranges from 30 - 50 years.^2^ The neurodegenerative disorder affects a variety of regions within the human brain such as globus pallidus, cerebral cortex, substantia nigra, subthalamic nuclei, hypothalamus and also cerebellum in the advanced stage of the disease but the most devastating effect of this disease is seen in striatum which is a part of basal ganglia.^3^ The symptoms ranges from uncontrolled spasmodic movements called Huntington’s chorea or simply put chorea to gait disturbances. Some people also experience behavior alterations, cognitive impairment, dysphagia in addition to the above stated symptoms.^4^ At present diagnosis of HD depends on the arrival of symptoms but by the time the disease has been detected, it has advanced to an irreversible stage. HD is a genetic disease and is diagnosed with the help of genetic screening test. Presently the treatment of HD is symptomatic which can delay the disease but cannot completely reverse the course of HD.^5^

Potential biomarkers for HD is the need of the hour. At present there are several biomarkers used for the detection of HD & they are -

i. Clinical biomarkers – determined using Anti saccade error rate^6^, Digitomotography^7^.
ii. Imaging – MRI^7^, MNI-659 PET^8^
iii. Immune biomarkers – IL8^9^
iv. Neurodegeneration biomarker – NfL^10^

Blood based biomarkers are going to play a key role in early detection of the disease. Blood based biomarkers are going to play a key role in the early detection of HD because the starting material (in this case blood) is easier to collect and requires less expertise for the collection procedure.

Currently there is lack of exact blood-based biomarker for HD. Besides early detection of HD, blood biomarkers open up new window for designing novel therapeutic drugs in the future which can reverse HD completely.

Blood based biomarkers can be used profiled using many ways such as RNA-seq and microarray data analysis. Differential gene expression analysis from blood samples of HD patients and normal controls can give us valuable information regarding biomarkers as well as the pathways associated with the disease. In one of the study using RNA-seq data it was found out that there were 5genes (PROK2, ZNF238, AQP9, CYSTM1 and ANXA3) that were significantly expressed more than other genes in HD samples.^11^

Microarray data were used for finding biomarker for disease like Alzheimer’s disease.^12^ Although gene expression profiling is an useful method but they have their own cons. Many incongruities occur during gene expression profiling which may occur due to designing of the experiment, storage method of blood samples and age of blood samples which directly effect the amount of RNA contained within the sample.^13^ So there is high chance that the genes which are expressed significantly in one study may not be expressed to that extent in other and therefore it is highly required that multiple datasets are studied at the same time which reduces this risk.

In the present study, three microarray datasets were obtained from gene expression omnibus database and are then analyzed for the purpose of finding a potential biomarker for HD.

## Methods

### Dataset Selection

Gene Expression Omnibus (GEO) by National Center for Biotechnology Information (NCBI) is an internationally acclaimed online database for high-throughput sequencing data, micro array, hybridization array data.^14^ GEO was searched for datasets in which keywords like “Huntington’s”, “blood”, “expression profiling by array”, “*Homo sapiens*” were present. There were three datasets which match the above criteria and their GEO accession numbers are GSE1751, GSE1767, GSE24250.^15,16^ All the datasets which are used in this study are available online and no actual human experiments were performed by the author. In this study total 32 patients samples and 34 normal control subjects were present. All the three studies used microarrays for data collection. Microarray is a chip based technology in which 1000s of nucleic acids are bound to a surface and are used to measure relative concentration of nucleic acid sequences in a mixture.^17^

**Table 1.**
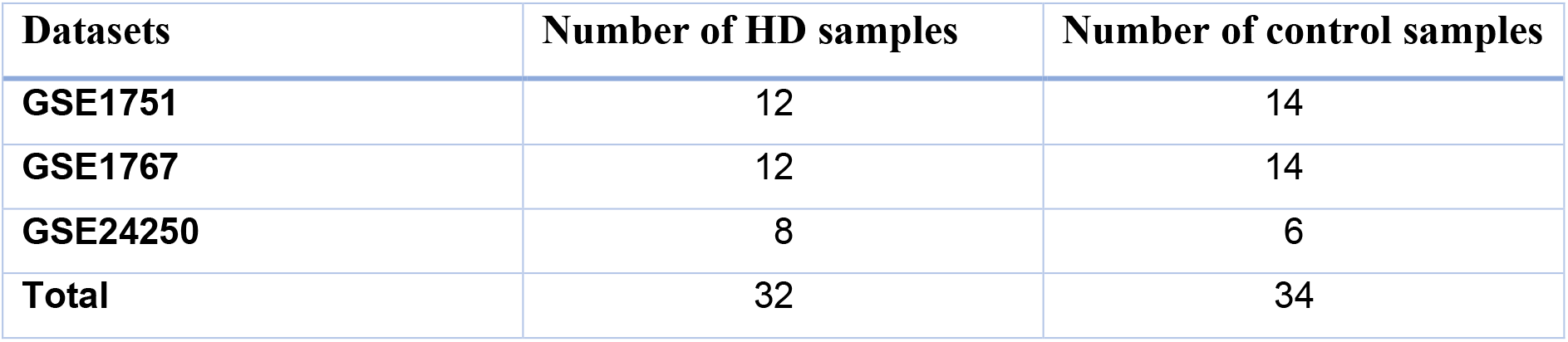
Dataset information for Microarray datasets obtained from GEO. There are in total 12 HD samples and 14 normal control samples in the dataset GSE1751, 12 HD samples and 14 normal control samples in the dataset GSE1767 and 8 HD samples and 6 normal control samples in the dataset GSE24250.

| Datasets | Number of HD samples | Number of control samples |
| --- | --- | --- |
| **GSE1751** | 12 | 14 |
| **GSE1767** | 12 | 14 |
| **GSE24250** | 8 | 6 |
| **Total** | 32 | 34 |

### Differential Gene Expression Analysis

The above 3 datasets were analyzed using GEO2R tool (available at http://www.ncbi.nlm.nih.gov/geo/geo2r/) which is provided by the Gene Expression Omnibus.GEO2R is an online software that processes gene expression values and outputs a table of two differentially expressed genes (DEGs) between two user defined groups (HD and control in this case). After running GEO2R, several thousand genes were found to be significant. p values were found out using ‘t’ test. Fold changes of each gene was calculated by the help of GEO2R tool. Fold change is represented by logFC values which tells us about the magnitude of gene expression change. So a value of 2.5 is 2 to the power 2.5 (2^2.5). This means that levels of gene expression for this gene are 5 times higher in patients than the normal subjects.

GEO2R tool was used for verifying a normal distribution of samples. No outliers were present in these datasets.

After obtaining the list of DEGs, they were processed in Excel. DEGs with p value greater than 0.05 were removed and the rest of the DEGs were sorted into two categories, which are overexpressed and those which are underexpressed. Some of the genes were overexpressed in HD while some are underexpressed. In this study only those genes were selected which have a fold change value greater than 1.2.

**Figure 1.**
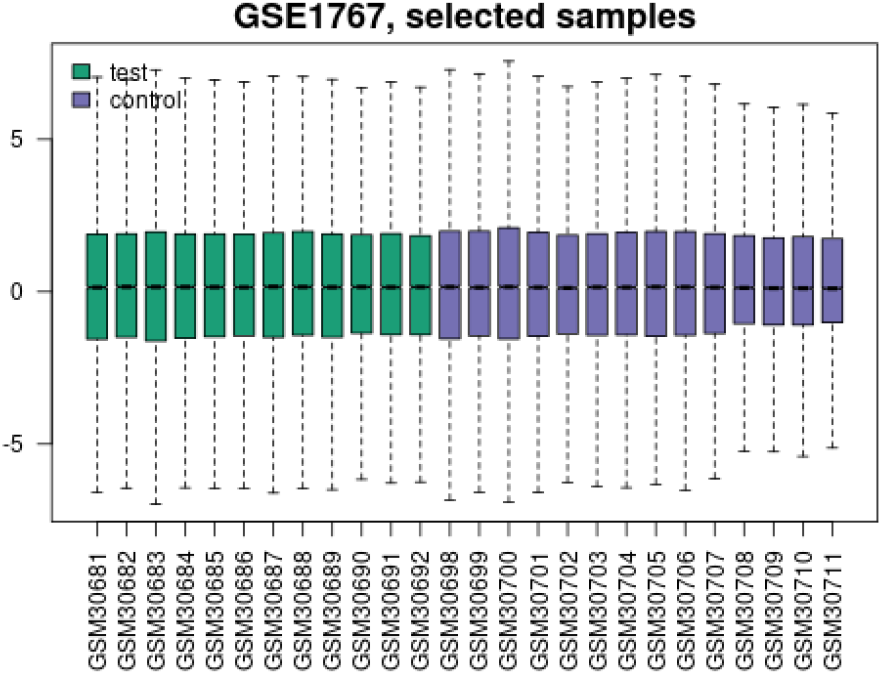
Gene expression value distribution for dataset GSE1767.There are no units for in y axis. Each box plot reperesents gene expression value of one patient sample.

**Figure 2.**
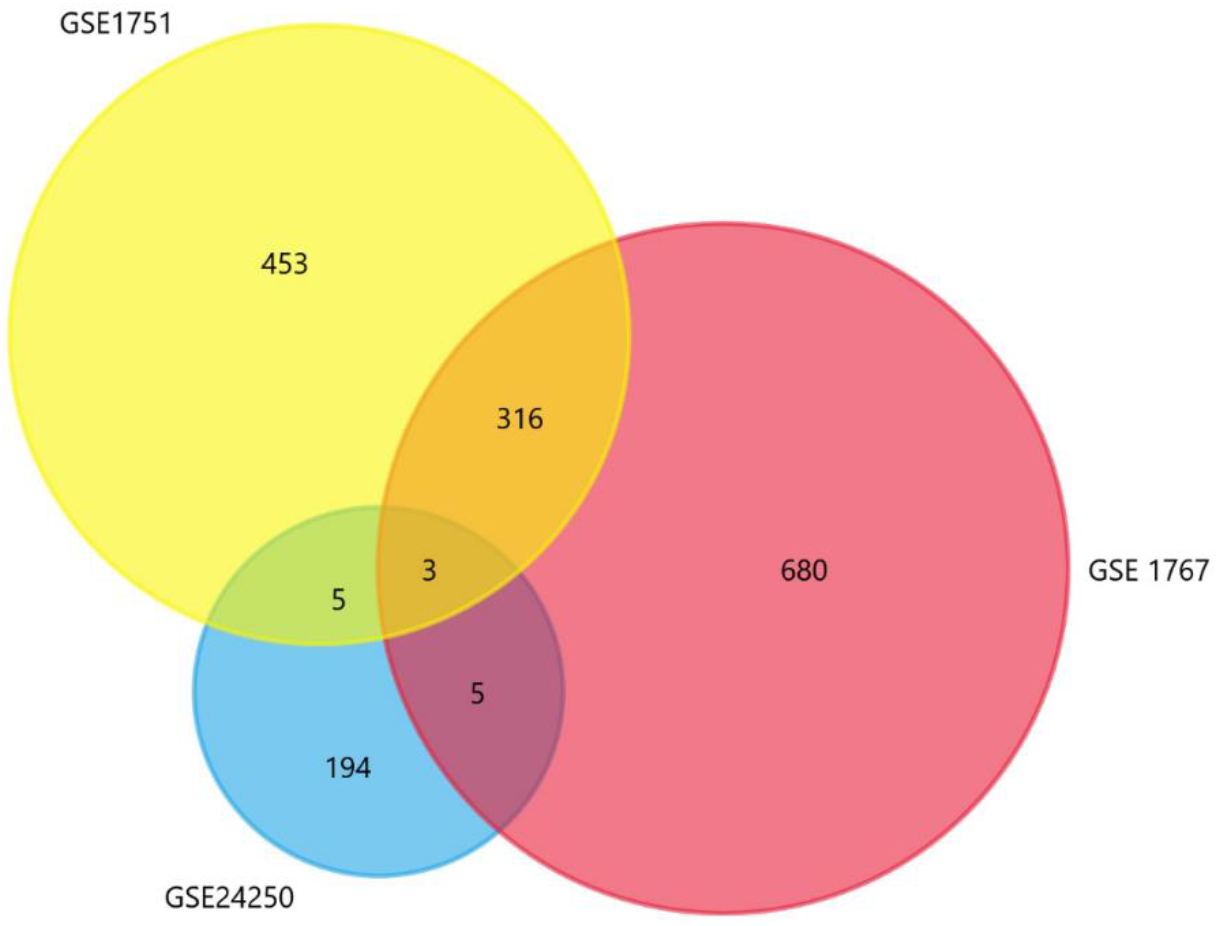
Venn diagram representing the DEGs in all the three datasets (GSE 1751, GSE 1767 and GSE 24250).

### Identification of Common Genes

Desktop based tool Fun Rich was used for creating Venn diagram in order to find out the common genes among all three datasets.^18^

### Network Analysis

The final list of DEGs were analyzed by the help of STRING (available at https://string-db.org/) which is used to construct a network of proteins which interact with each other. STRING is a search tool for interacting proteins and also calculates statistically significant Gene Ontology processes and pathways.^19^ A tab separated value file was obtained from STRING and was then imported to Cytoscape. Cytoscape is an open source desktop based software used for integrating biomolecular interaction networks with the high-throughput expression data.^20^ Network analyzer, a Cytoscape plug-in CytoHubba was used for the identification of hub genes.

### Enrichment Analysis of DEGs

Gene enrichment analysis was performed for DEGs using the online software called Enrichr (available at https://maayanlab.cloud/Enrichr/) for Gene Ontology(GO) function and also for KEGG pathway enrichment. GO terms were further subdivided into several terms such as Biological Process, Molecular Function and Cellular Component.

### Results

Three gene expression profiles (GSE 1751, GSE 1767 and GSE 24250) were analyzed with the help of GEO2R tool and the list of DEGs were downloaded. p value was set at 0.05 and a log FC value of 1.2 was set as the cut off criteria. There were 903 genes in the dataset GSE 1751, 1168 genes in the GSE 1767 and 229 genes passed the above criteria in the dataset GSE 24250. Venn analysis was performed using the software Fun Rich and the DEGs were identified. 329 genes were found to be significantly differentially expressed among all the three above mentioned datasets. In these datasets there were three genes which are differentially expressed in all the datasets and the name of those genes are FNDC3A, BCLAF1 and ALCAM. FNDC3A (Fibronectin type-III domain-containing protein 3A) is a protein coding gene which produces a transmembrane protein present in human odontoblast and dental pulp nerves and to a much lesser extent in human brain, kidney, liver and lungs.^21^ Not much is known with respect to the function of this protein. BCLAF1 (Bcl-2-associated transcription factor 1) is a transcription factor which is associated with several biological process such as apoptosis, negative as well as positive regulation of transcription, regulation of DNA template regulated transcription in response to stress.^22,23^ ALCAM (Activated Leukocyte Cell Adhesion Molecule) encodes a protein called CD166 which is a transmembrane glycoprotein which is a cell adhesion molecule which mediates heterotypic cell adhesion, T cell activation, normal cell enfragment in the bone marrow.^24,25^

The 329 genes which passed the above cut-off criteria were submitted to the STRING tool. The disconnected genes were hidden and medium confidence option was selected in the minimum required interaction score option. Total of 326 nodes and 964 edges are present in the network(Figure 3). The online tool Enrichr was used for performing GO and KEGG pathway enrichment analysis. After submitting the list of DEGs the the ontology option in the Enrichr gave several options such as GO Biological Process (GO-BP), GO Molecular Function (GO-MF), GO Cellular Component (GO-CC). GO analysis showed that the DEGs were enriched in GO-BP such as negative regulation of programmed cell death, regulation of transcription from RNA polymeraseII promoter in response to hypoxia, negative regulation of apoptosis, toll like receptor signalling pathway, MYD88-independent toll like receptor signalling pathway,regulation of apoptotic process,regulation of cytokine mediated signaling pathway, activation of cysteine-type endopeptidase activity invoved in apoptotic signaling pathway(Fig 4A). GO-MF analysis showed that DEGs were specially enriched in cadherin bingding, RNA binding, kinase binding, protein heterodimerization activity, cysteine type endopeptidase inhibitor activity involved in apoptostic process, protein kinase binding, nucleoside triphosphatse activity, metal ion transmembrane activity, purine ribonucleoside triphosphate binding and annealing acitivity(Fig 4B). For GO-CF analysis, the DEGs were mainly enriched in nuclear body, nuclear speck, nucleolus, cytosolic part, cytosolic ribosome, cyclin/CDK positive transcription elongation factor complex, chromosome, cytosolic small ribosomal subunit, ribosome(Fig 4C). The KEGG pathway analysis showed that DEG are enriched in pathways which are associated with NOD-like receptor signalling pathway, Kaposi sarcoma-associated herpesvirus infection, Small cell lung cancer,Nf-kappa B signalling pathway, Apoptosis, paltelet activation, phosphatidyl inositol signalling system, several pathways in cancer and toxoplasmosis and also Human cytomegalovirus infection(Fig 4D). For finding out the hub gene, the genes with degree greater than equal to 15 were selected from the list of DEGs and was submitted to the Cytoscape offline tool.In the cytohubba plug-in it was found out that Phosphatse and tensin homolog has the highest connectivity with other genes (PTEN, degree = 18) followed by Interleukin 8 (CXCL8, degree = 18), 26s proteasome regulatory subunit 10B (PSMC6, degree = 15), Caspase 3 (CASP3, degree = 15), Ubiquitin Conjugating Enzyme E2D1 (UBE2D1, degree = 14), CD27 (CD27, degree = 14), Splicing factor 3b subunit 1 (SF3B1, degree = 14), Eukaryotic translation initiation factor 4E (EIF4E, degree = 14), Baculoviral IAP repeat-containing protein 2 (BIRC2, degree = 14) (Figure 5). These above stated genes are upregulated in HD.

**Figure 3.**
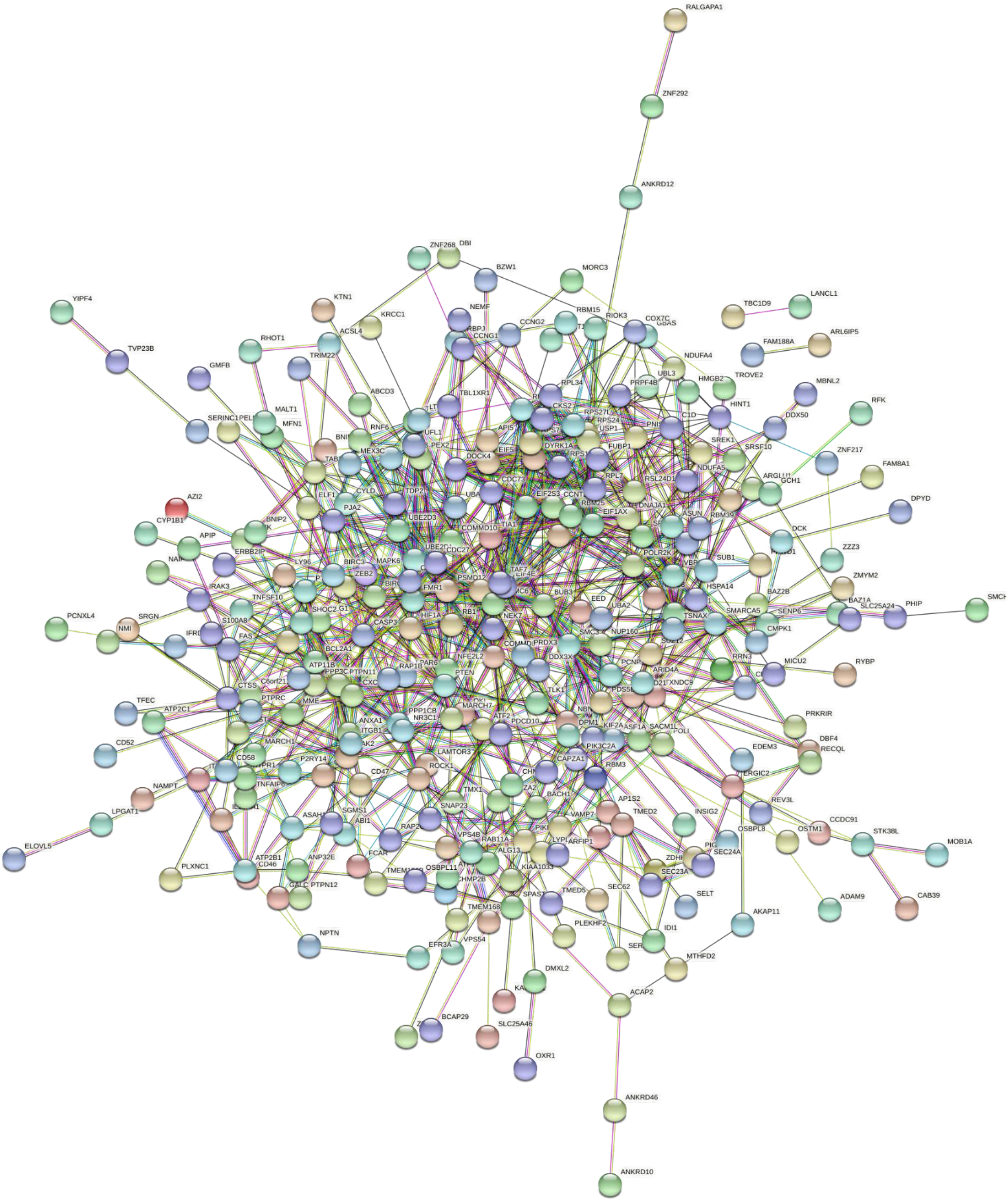
STRING protein protein interaction network. In this network there are 326 nodes and 964 edges. Lines represents interaction and the circles represent genes. The colour of the lines represents the type of interaction.

**Figure 4.**
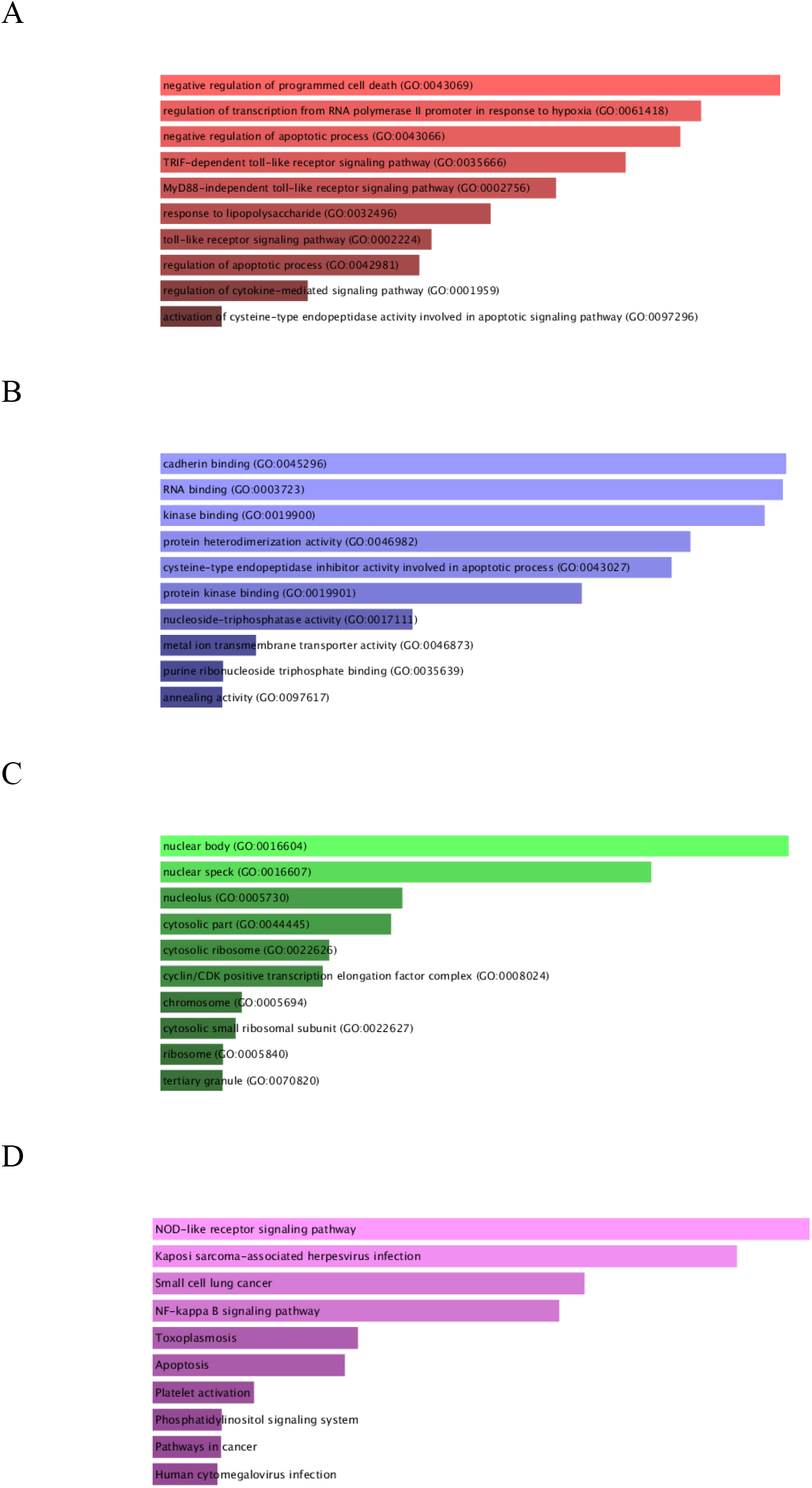
GO and KEGG pathway enrichment analysis using Enrichr. A-Top 10 enriched biological processes in DEGs. The x axis represents number of genes and the y axis represents the biological process. B-Top 10 enriched molecular function in DEGs. The x axis represents number of genes and the y axis represents molecular function. C-Top 10 enriched cellular components in DEGs. The x axis represents the number of genes and the y axis represents cellular components. D – Top 10 enriched KEGG pathways for DEGs. The x axis represents the number of genes and the y axis represents the KEGG pathway names.

**Figure 5.**
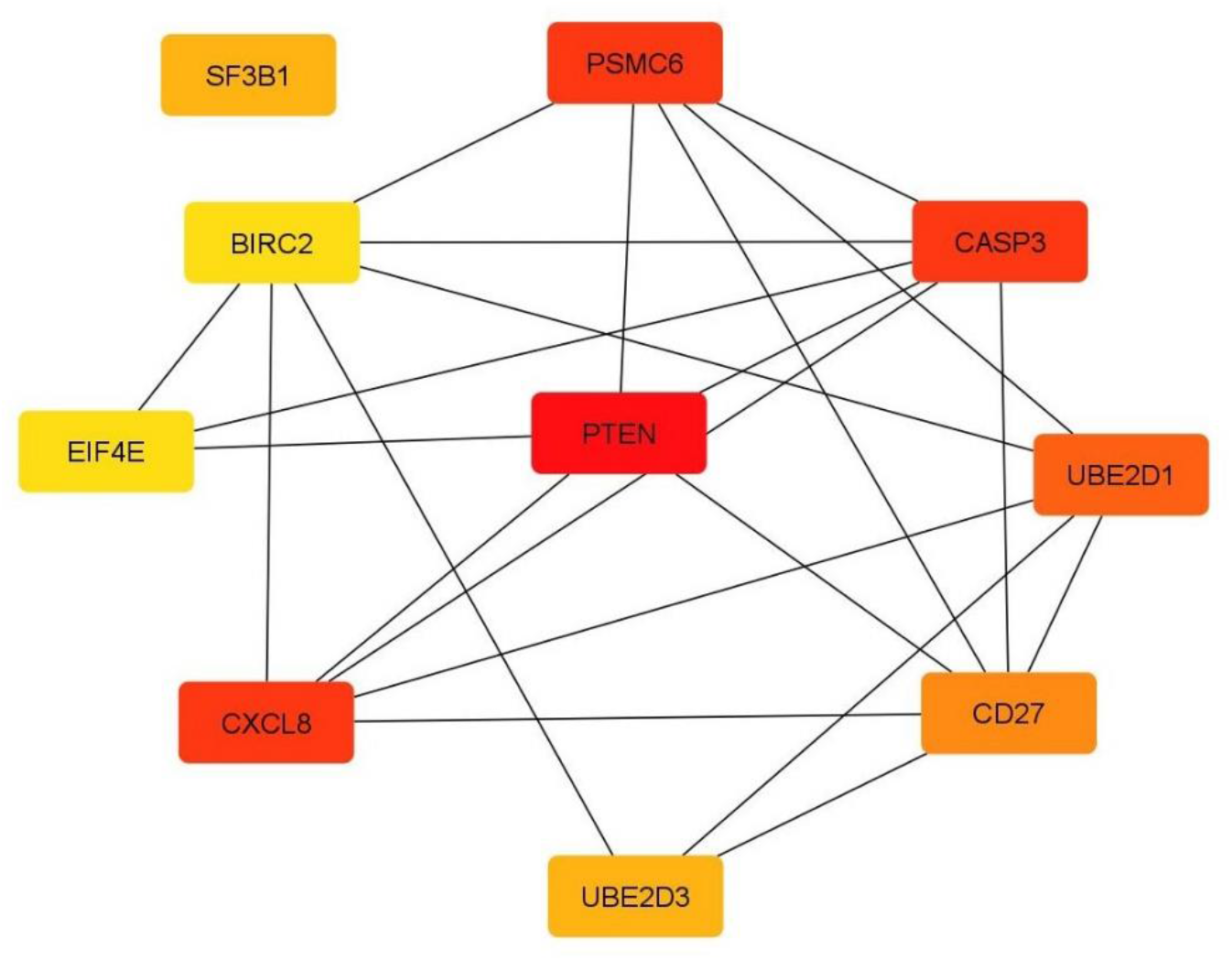
Network of top ten hub genes using Cytoscape software. The colour represents the degree of connectivity, red colour represents the highest degree, the orange colour represent intermediate degree and the yellow represent the lowest degree.

## Discussion

Huntington’s disease is a neurodegenerative disease associated with many cellular changes and in this study some key genes were identified which were earlier not reported. Genes such as CXCL8, PSMC6, UBE2D1, UBE2D1, CD27, UBE2D3, SF3B1were previously not known to regulate HD. Similarly there were some genes which are already known to regulate HD such as CASP3, EIF4E, BIRC2.^26–28^ Upregulation PTEN occurs in several neurodegenerative diseases.^29^ Several pathways, biological functions associated with the above mentioned upregulated genes were also enriched in the KEGG and GO enrichment analysis. However the three genes namely FNDC3A, BCLAF1 and ALCAM which were present in all the three datasets were not the hub genes.

Limitations of this study includes that there was no data normalization in this study and no distinction has been made between male and female patients as well as normal controls where there can be significant difference in differentially expressed genes with respect to gender. Therefore further study on this subject is required so that new potential blood based biomarkers of HD can be found.

